# Dormancy dynamics and dispersal contribute to soil microbiome resilience

**DOI:** 10.1101/730283

**Authors:** Jackson W Sorensen, Ashley Shade

## Abstract

In disturbance ecology, stability is composed of resistance to change and resilience towards recovery after the disturbance subsides. Two key microbial mechanisms that can support microbiome stability include dormancy and dispersal. Specifically, microbial populations that are sensitive to disturbance can be re-seeded by local dormant pools of viable and reactivated cells, or by immigrants dispersed from regional metacommunities. However, it is difficult to quantify the contributions of these mechanisms to stability without, first, distinguishing the active from inactive membership, and, second, distinguishing the populations recovered by local resuscitation from those recovered by dispersed immigrants. Here, we investigate the contributions of dormancy dynamics (activation and inactivation), and dispersal to soil microbial community resistance and resilience. We designed a replicated, 45-week time-series experiment to quantify the responses of the active soil microbial community to a thermal press disturbance, including control mesocosms, disturbed mesocosms without dispersal, and disturbed mesocosms with dispersal after the release of the stressor. Communities were sensitive within one week of warming. Though the disturbed mesocosms did not fully recover within 29 weeks, resuscitation of thermotolerant taxa was key for community transition during the press, and both resuscitation of opportunistic taxa and immigration contributed to community resilience. Also, mesocosms with dispersal were more resilient than mesocosms without. This work advances the mechanistic understanding of how microbiomes respond to disturbances in their environment.

## Introduction

Ongoing changes to Earth’s climate are projected to alter disturbance regimes and to pervasively expose ecosystems to stressors like elevated atmospheric greenhouse gases and increased temperatures[1]. Microbial communities, or *microbiomes*, provide vital ecosystem functions and are key players in determining ecosystem responses to environmental changes[2,3]. Understanding the mechanisms that underpin microbiome responses to environmental disturbances will support efforts to predict, and, potentially, manage, microbiomes toward stable functions within their ecosystems.

In disturbance ecology, stability refers to consistent properties in the face of a stressor [4]. Here, we apply terms from disturbance ecology as they have been adopted in microbial ecology[5,6]. Stability includes components of both resistance and resilience. Resistance is the capacity of a system to withstand change in the face of a stressor, and its inverse is sensitivity. Resilience is the rate of return following a disturbance. Recovery is when a system fully returns to its pre-disturbance state, and an alternative stable state is when the system does not return but rather assumes a different state. Together, resistance, resilience and recovery are the major quantifiable components of stability, and they can be calculated from community measurements of alpha diversity, beta diversity, or function[6,7].

There are two related microbial mechanisms that support population persistence in the face of disturbance, and therefore contribute to community resistance, resilience, and recovery. One mechanism is microbial dispersal, as successful immigrants can support resilience and recovery of sensitive populations. Across an interconnected landscape, microbial metacommunities are linked via dispersal, and so immigrants originate from the regional species pool [8–11]. A second important but less-considered mechanism is microbial dormancy dynamics [12,13]. Dormancy dynamics include initiation and resuscitation. Initiation into dormancy can support local survival of populations sensitive to the disturbance, and therefore support community resistance. Resuscitation from dormancy can support resilience and recovery by re-seeding sensitive populations from the local dormant pool. Thus, while both dispersal and resuscitation can support microbiome stability, dispersed immigrants originate regionally while resuscitated members originate locally. After a disturbance, if sensitive populations are not repopulated via immigration or resuscitation, they will become locally extinct and contribute to standing necromass (aka relic DNA, [14]).

We designed a replicated time-series experiment to quantify the contributions of dormancy dynamics and dispersal to the response of a soil microbiome to a thermal press disturbance. We targeted a soil microbiome because terrestrial microbiomes are front-line responders to climate change and sequesters of carbon [2,3], and therefore an important constituent to understand for predicting ecosystem outcomes to environmental change. Also, soils harbor the highest known microbial diversity [15–17] and present a maximum challenge in deciphering microbiome responses to disturbance. Furthermore, a majority of the microbial cells or richness in soil is reportedly dormant [12,18], reportedly as high as 80%, representing a considerable pool of microbial functional potential. Finally, across heterogeneous soils, an average of 40% of the microbiome DNA was necromass that existed extracellularly[14]. This suggests that DNA-based methods of determining microbiome dynamics include both inactive and necromass reservoirs, and that there is need for increased precision to move forward to quantify mechanisms underpinning microbiome stability.

The mesocosm experiment reported here follows our prior field work in Centralia, Pennsylvania [19–23]. Centralia is the site of an underground coal seam fire that ignited in 1962 and advances 5-7 my^-1^ along the coal seams[24,25]. The coal seams are highly variable in depth, but average 70 m below the surface[24], so as the fire advances underground it warms the overlying surface soils to mesothermal to thermal conditions. After the fire advances, previously warmed soils cool to ambient temperatures. In the field, we observed that previously warmed soils recovered towards reference soils in bacterial and archaeal community structure, with the exception of a slightly increased selection for Acidobacteria in the recovered soils (attributable to lower soil pH after coal combustion,[19]). However, during fire impact, there was high divergence among soil communities, and we hypothesized that differences in dormancy dynamics (e.g., different members resuscitating and initiating priority effects during the stress) may explain the divergences. In this experiment, we aimed to control dispersal, and also to quantify activity dynamics and determine their consistency.

## Materials and Methods

### Soil collection, mesocosm design, and soil sampling

Eight kg of soil was collected in Whirlpack bags from the top ten centimeters of a reference site in Centralia, PA (site C08, 40 48.084N 076 20.765W) on March 31^st^, 2018. The site is temperate with the following chemical-physical properties: Organic Matter 4.8%; Nitrate 7.9 ppm; Ammonium 20.5 ppm; pH 5; Sulfur 19 ppm; Potassium 69 ppm; Calcium 490 ppm, Magnesium 59 ppm; Iron 110 ppm, and Phosphorus 395 ppm. The ambient soil temperature when collected was 4°C. The sample was stored at 4°C until the experiment was initiated. Soil was sieved through a 4mm mesh, homogenized, and ∼300 g were dispensed into 15 autoclaved quart-sized glass canning jars that were used as mesocosms (Ball). The homogenized soil sample intentionally was used in all 15 mesocosms to assess the reproducibility of community temporal dynamics starting from the same soil source. Percent soil moisture was determined using by massing and drying. Each mesocosm was massed weekly to assess evaporation and any loss of water mass was replaced with sterile water to maintain percent soil moisture throughout the experiment. Sterile metal canning lids were secured loosely to prevent anaerobiosis. All set-up and manipulation of the mesocosms was performed in a Biosafety Level 2 cabinet (ThermoScientific 1300 Series A2) and we used aseptic technique.

Mesocosms first were acclimated at 14°C to mimic the ambient soil temperature at the typical time of fall soil collection and to coordinate with the field study [19]. Acclimation proceeded for four weeks in a cooling incubator (Fischer Scientific Isotemp), and then soils were divided into three treatment groups (**Figure 1**). Six control mesocosms were maintained at 14°C for the duration of the experiment. Nine disturbance mesocosms were subjected to a 12-week disturbance regime to simulate a press thermal disturbance. First, the temperature was gradually increased to 60°C over two weeks (increase period). Second, the temperature was maintained at 60°C 8 weeks. Sixty degrees was chosen because it was close to the observed maximum thermal temperature that we have measured in surface soils impacted by the Centralia coal seam fire [19]. Next, the temperature was gradually decreased to 14°C over two weeks. Finally, the mesocosms were maintained at 14°C for four weeks until the penultimate sampling. From the nine disturbed mesocosms, four were randomly selected for the dispersal treatment. These four disturbed mesocosms received a dispersal event one week after the temperature was recovered to 14°C after the thermal disturbance. Each was inoculated with 0.5 mL of a 10% weight by volume soil slurry made from a composite soil sample from the six control mesocosms. We used soil from the control mesocosms to simulate dispersal from similar, adjacent soils to repopulate disturbed communities. Finally, all mesocosms were left undisturbed at 14°C for another 25 weeks prior to the final 45-week sampling. During the final 25 week incubation, percent moisture was not monitored.

**Figure 1.**
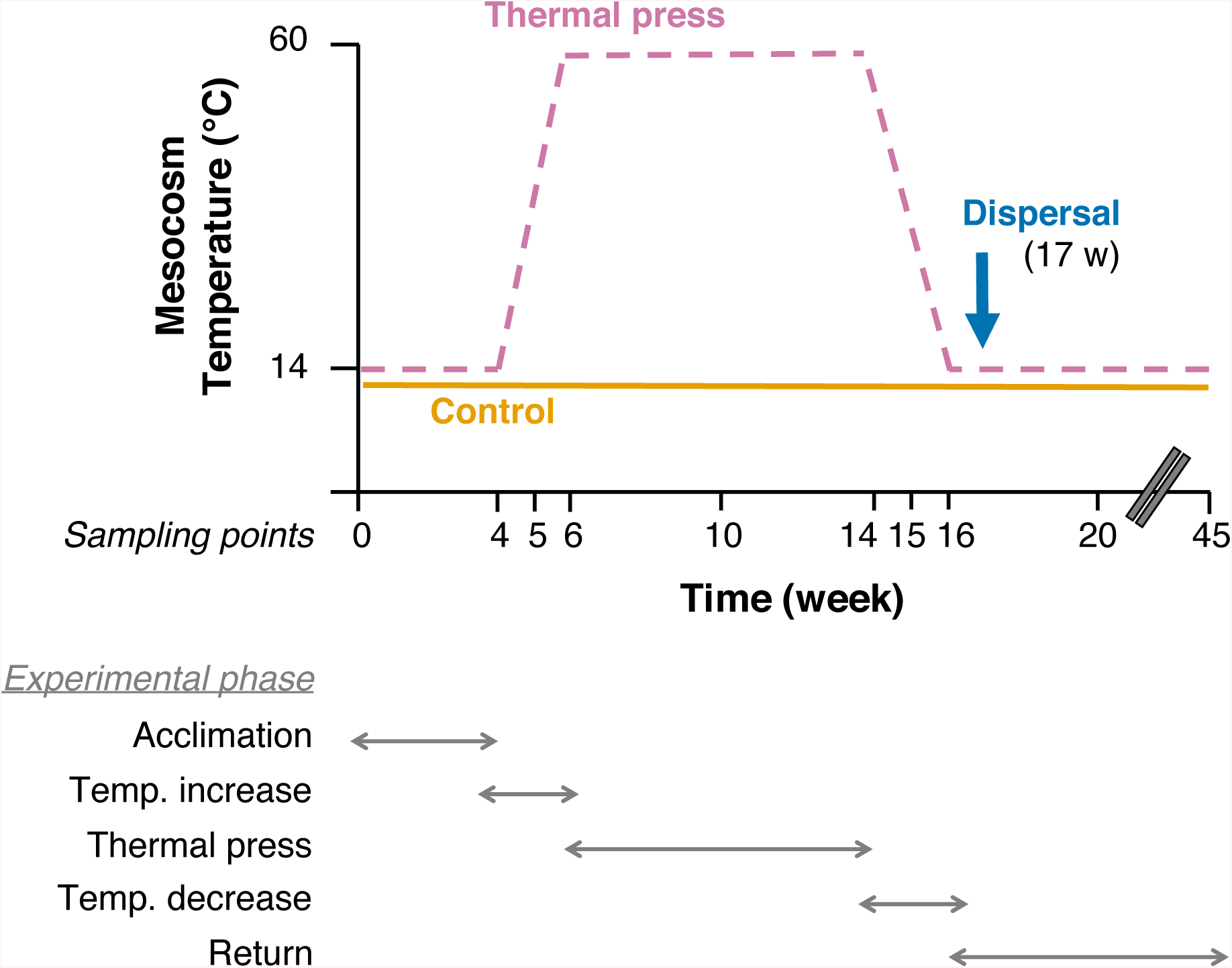
Experimental design of the study. Reference temperate soil (0-20 cm depth from surface) was homogenized and divided among fifteen 1 L glass mesocosms that were maintained at ambient moisture through the experiment. Control mesocosms (solid gold line, n = 6) were maintained at 14°C, which was ambient soil temperature at the time of collection. Disturbance mesocosms (dashed pink line, n = 9) were acclimated for four weeks at 14°C, increased to 60°C over two weeks, maintained at 60°C as a thermal press disturbance for eight weeks, then decreased back to 14°C over two weeks, and finally maintained for a total of 45 weeks. Four of the disturbance mesocosms received a dispersal event (homogenized soil slurry from Control mesocosms, see methods) at week 17, after the thermal press was released. Note the break in the x-axis time scale between weeks 20 and 45.

Mesocosms were non-destructively sampled after 4, 5, 6, 10, 14, 15, 16, and 20 weeks of incubation. At each time point, approximately 15 g soil was removed from a mesocosm, of which ∼13 g was flash-frozen in liquid nitrogen for RNA preservation and stored at −80°C until RNA/DNA co-extraction.

### RNA/DNA co-extraction

To obtain RNA and DNA from the same cell pool, we minimally modified a manual coextraction protocol originally published by [26]. For each sample, 0.5 g of flash-frozen soil was added to Qiagen PowerBead Tubes containing 0.70 mm garnet beads. Next, 500 uL of a 5% CTAB/Phosphate buffer and 500 uL of phenol:chloroform:isoamyl alcohol were added to each PowerBead tube. Cells were then lysed using a Model 607 MiniBeadBeater-16 (BioSpec Products Inc.) for 30 seconds, followed by a 10 min centrifugation at 10,000 x g and 4°C. The top aqueous layer was transferred to a fresh tube and 500 uL chloroform:isoamyl alcohol was added. The tubes were inverted several times to form an emulsion before a five minute centrifugation at 16,000 x g and 4°C. The top aqueous layer was transferred to a clean 1.5 mL centrifuge tube. Nucleic acids were precipitated by adding two volumes of a 30% PEG6000 1.6M NaCL solution, inverting several times to mix, and incubating on ice for two hours. After incubation, nucleic acids were pelleted by a 20 min centrifugation at 16,000 x g and 4°C. The supernatant was removed from each tube and one mL of ice-cold ethanol was added to the pelleted nucleic acids. Tubes were centrifuged for 15 min at 16,000 x g and 4°C, and the ethanol supernatant was removed. Pelleted nucleic acids were left to air dry before resuspending in 30 uL of sterile DEPC-treated water.

To purify the RNA, co-extracted nucleic acids were diluted 1:100 before treatment with Ambion Turbo DNA-free DNase kit, using the robust treatment option in the manufacturer’s instructions. Extracted nucleic acids were mixed with 0.1 volumes of the 10X Turbo DNase Buffer and three uL of TURBO Dnase enzyme (six units total) and incubated at 37°C for 30 min. After incubation, 0.2 volumes of DNase inactivation reagent was added and incubated for five minutes at room temperature before a five min centrifugation at 2,000 x g and room temperature. The treated supernatant was removed and used as the template for Reverse Transcription. RNA purity was assessed by PCR (see below for details) and showed no amplification. Reverse transcription was performed with random hexamers using the SuperScript III First-Strand Synthesis System for RT-PCR(Invitrogen) per manufacturer’s instructions.

PCR of cDNA and no-RT controls was performed using the Earth Microbiome Project 16S V4 primers(515F 5’-GTGCCAGCMGCCGCGGTAA-3’, 806R 5’-GGACTACHVGGGTWTCTAAT-3’) [15,27]. Temperature cycling was as follows: 94°C for four minutes followed by 30 cycles of 94°C for 45 seconds, 50°C for 60 seconds and 72°C for 90 seconds followed by a final elongation step at 72C for 10 minutes. Products were visualized using gel electrophoresis.

### 16S rRNA and 16S rRNA gene sequencing and processing

Here, for simplicity we use “microbiome” to refer to the bacterial and archaeal community members captured by amplifying and Illumina sequencing of the 16S ribosomal RNA and DNA (rRNA gene). Library preparation and sequencing was performed by the Michigan State University Genomics Core Research Facility. A single library was prepped using the method in Kozich et al (2013) [28]. PCR products were normalized using Invitrogen SequalPrep DNA Normalization Plates. This library was loaded onto 4 separate Illumina MiSeq V2 Standard flow cells and sequenced using 250bp paired end format with a MiSeq V2 500 cycle reagent cartridge. Base calling was performed by the Illumina Real Time Analysis (RTA) V1.18.54.

All samples were first checked for any contaminating primer sequences using cutadapt[29], before being processed together using the USEARCH pipeline[30,31]. Briefly, paired end reads were merged using -fastq_mergepairs and then dereplicated using -fastx_uniques. Reads were clustered *de novo* at 97% identity and then the original merged reads were mapped to the representative sequences of each cluster. Each OTU was classified using SINTAX[32] and with the Silva database (version 123, [33]).

### Designating Total and Active Communities

Each RNA and DNA sample was rarefied to 50,000 reads in R using the vegan package version 2.5-4 [34] discarding any samples which did not contain sufficient reads. Samples for which either the RNA or DNA did not have 50,000 reads were omitted from the analysis presented here (12 out of 135 in total). The Total community was defined as the community recovered in the DNA reads. The Active community was defined per sample, using the DNA read numbers of those taxa that had 16S rRNA:rRNA gene ratio was >1 in each sample[35]. Consequently, while every sample was initially rarefied to 50,000 reads, each sample’s active community varied slightly in total reads.

### Quantitative PCR (qPCR)

qPCR was performed on the V4 region of the 16S rRNA gene and conducted in a BioRad CFX qPCR machine using the Absolute QPCR Mix, SYBR Green, no ROX (Thermo Scientific). Each reaction contained 12.5ul of the 2X Absolute QPCR Mix, 1.25 ul each of 10uM primers 515F and 806R, 3uL of template DNA and 2uL of PCR grade water. Temperature cycling conditions were as follows: 15 minutes at 95°C, followed by 39 cycles of 94°C for 45 seconds, 50°C for 60 seconds, and 72°C for 90 seconds, followed by a final elongation step at 72°C for 10 minutes. Fluorescence was measured in each well at the end of every cycle. Extracted gDNA from *E. coli* MG1655 was used for the standard curve, which was run in triplicate with every plate. Samples were run in duplicate across different plates and those that amplified after the lowest point of the standard curve (27 copies per reaction) were treated as zeroes. No template controls were included in every qPCR plate and they never amplified. Amplification specificity was assessed by melt curve (60°C to 95°C, 0.5°C increments).

### Ecological statistics

Ecological analyses were performed in R[36]. The adonis function in the vegan package was used to perform PERMANOVAs[37], and the betadisper function was used to quantify beta dispersion[38] with Tukey’s Honestly Significant Difference post-hoc test. Pairwise tests for alpha diversity, community size, and resilience values were performed using the Kruskal-Wallis test, with Dunn’s post-hoc correction for multiple comparisons when needed. Principal coordinates analysis was used for ordination of pairwise sample differences based on Bray-Curtis dissimilarity. Procrustes superimposition (PROTEST) was performed using the procrustes function in the vegan package and a false discovery rate adjustment was used for multiple tests. Data visualizations were performed using ggplot2[39]. Heatmaps were made using the heatmap.2 function in the gplots package[40].

Contributions of responsive and immigrant taxa to beta diversity were calculated as the Bray-Curtis dissimilarity attributed to the responsive taxa subset and divided by the total Bray-Curtis dissimilarity, both calculated from the Total (DNA) community, as done previously to assess the contributions of conditionally rare taxa to beta diversity [41]. Responsive taxa were those that changed in activity between weeks 16, 20, and 45 by their 16S rRNA:rRNA gene, either from < 1 to > 1 or > 1 to < 1. Immigrant taxa were undetected in all disturbed mesocosms at week 16, but detected in Disturbance + Immigration mesocosms at either week 20 or week 45 while remaining undetected in the Disturbance mesocosms.

### Data availability and code

Sequence workflows, OTU tables, and statistical workflows to reproduce the analyses described here are available on GitHub (https://github.com/ShadeLab/PAPER_Sorensen_InPrep_Mesocosms). All raw sequence data are deposited in the NCBI Short Read Archive under BioProject PRJNA559185.

## Results

### Sequencing summary

In total, we sequenced 135 pairs of samples (cDNA and DNA) across nine timepoints and 15 mesocosms. We rarefied all samples to 50,000 reads, and removed those samples with fewer than 50,000 reads. This resulted in the removal of 12 samples and left 53 Control, 36 Disturbance, and 34 Disturbance + Immigration pairs of samples. After rarefaction, sample richness ranged from 84 to 4,108, with 16,854 total OTUs observed, inclusive of both DNA and RNA datasets.

### Overarching responses to the thermal press disturbance

Total community richness responded consistently and as expected to the thermal press disturbance. There was a notable bottle effect of maintaining field soil in mesocosms, indicated by the gradual decrease in richness over time in the Control treatment (**Figure 2AB**). In the Disturbance treatment, there was a modest but statistically supported decrease in richness one week after warming from 14°C to 37 °C (week 5 all Disturbance v. Control comparison, Kruskal-Wallis test, p = 0.003), and then a more substantial decrease after warming to 60°C at week 6 (Kruskal-Wallis test, p = 0.002). Community size was estimated using copies of the 16S rRNA gene measured with qPCR (**Figure 3**). Disturbance community size decreased over weeks four to seven and then maintained at a median of 1.03 x 10^7^ rRNA gene copies per g soil. Control communities decreased until week seven (bottle effect) and then increased rapidly by week ten and generally stabilized at median of 2.98 × 10^8^ 16S rRNA gene copies/g soil (**Figure 3A**). Together, these results show that the warming treatment acted as an environmental filter, resulting either in death or population decreases past the limits of detection for taxa that were otherwise fit in temperate conditions. Furthermore, there was a small but appreciable increase in richness after the dispersal event in the Disturbance + Immigration treatment, relative to the Disturbance treatment (Kruskal – Wallis test p= 0.088 at week 20, and p = 0.168 at week 45), and this increase was also observed for community size, which approaches recovery towards the control (Kruskal – Wallis test Control vs Disturbance + Immigration p=0.11, Control vs Disturbance p=0.0004, Disturbance vs Disturbance + Immigration p=0.013) (**Figure 3B**). This suggests that the dispersal treatment was effective in promoting recovery of richness and community size. However, warmed mesocosms did not completely recover richness to the level of the ambient Controls, even by week 45 (**Figure 2B**). Evenness followed the same overarching patterns as richness (**Figure 2CD**).

**Figure 2.**
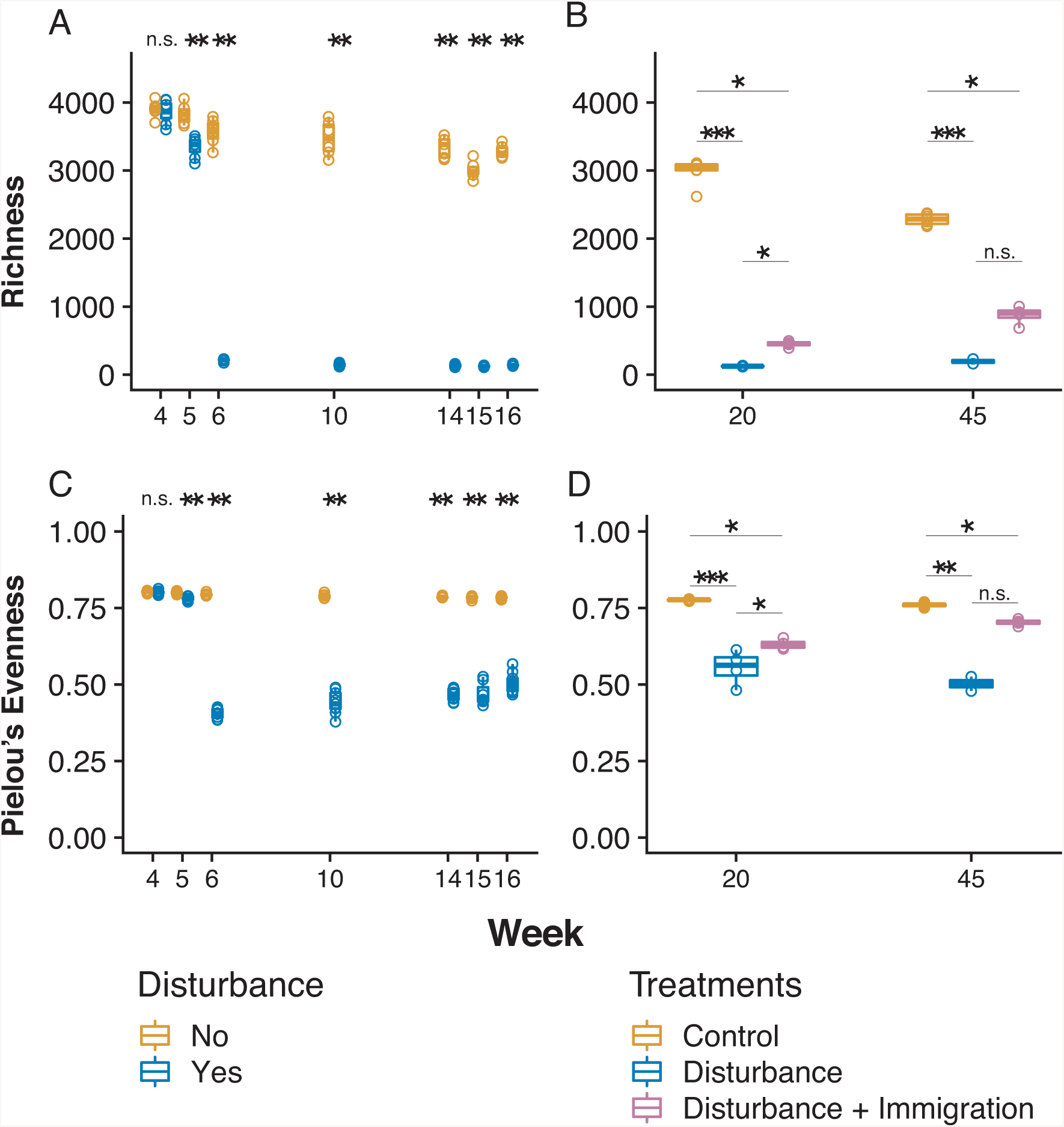
Changes in alpha diversity over the disturbance experiment. Alpha diversity was assessed using operational taxonomic units clustered at 97% sequence identity, after 16S rRNA gene sequencing and rarefaction to 50,000 sequences per sample. (A) Changes in the observed no. OTUs (richness) in Control (gold) and Disturbance (blue) mesocosms over the thermal press (weeks 4-16). (B) Changes in richness in Control, Disturbance, and Disturbance + Immigration (pink) mesocosms over the recovery period, weeks 20-45. The Disturbance + Immigration mesocosms received a dispersal event at week 17. (C) Changes in evenness over weeks 4-16. (D) Changes in evenness over weeks 20-45. Asterisks indicate significant differences by a Kruskal Wallis test (n.s = not significant; * p<0.1, ** p<0.01, *** p<0.001, with a Dunn correction for multiple comparisons in B and D).

**Figure 3.**
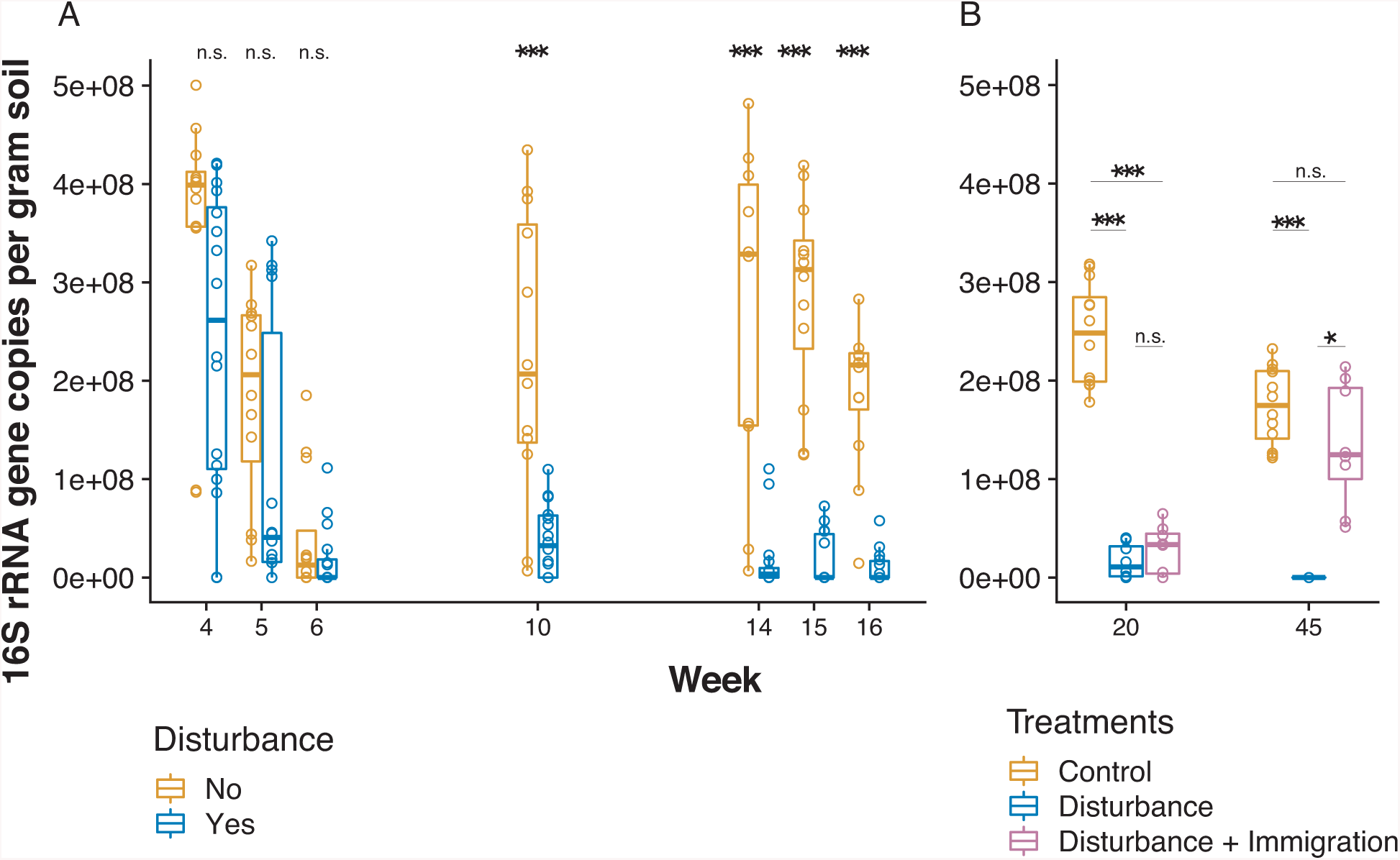
Changes in community size over the disturbance experiment. Community size was estimated using qPCR of the 16S rRNA gene and standardized per gram of soil from which nucleic acids were extracted. (A) Changes in the 16S rRNA gene copies in Control (gold) and Disturbance (blue) mesocosms over the thermal press (weeks 4-16). (B) Changes in the 16S rRNA gene copies in Control, Disturbance and Disturbance + Immigration (pink) mesocosms over the recovery period, weeks 20-45. The Disturbance + Immigration mesocosms received a dispersal event at week 17. Asterisks indicate significant differences by a Kruskal Wallis test (n.s. = not significant, * p<0.1, ** p<0.01, *** p<0.001, with a Dunn correction for multiple comparisons in B).

We compared community structure across treatments for the Total community dataset, rRNA gene; 14,159 OTUs) and the Active dataset (rRNA:rRNA gene > 1; 6,693 = OTUs). There were clear and consistent shifts in beta diversity in the Disturbance mesocosms, as well as high reproducibility among replicates for all treatments (**Figure 4, Figure 5**). Over the experiment, Disturbance mesocosms had distinct community structures than Control (Disturbance v. Control PERMANOVA PsuedoF = 63.87, Rsqr = 0.345, p=0.001 for Total communities, and PsuedoF=35.97, Rsqr=0.229, p=0.001 for Active communities). Control communities were relatively stable over the study, while Disturbance communities changed directionally, and were significantly different from Control communities after a single week of warming (week 5 Control vs Disturbed PERMANOVA PsuedoF = 3.06, Rsqr= 0.218, p=0.001 for Total community and PsuedoF= 2.88, Rsqr=0.208, p=0.001 for Active community). Disturbance communities continued to shift with temperature during the course of the experiment, and then shifted back towards the Control after the stressor was released. Though no Disturbance mesocosms fully recovered to overlap with the Control communities, the mesocosms with dispersal achieved more complete recovery than those without. Total communities and Active communities were synchronous in their temporal trajectories (Mantel R =0.943, p = 0.001 on 999 permutations; Protest Sum of Squares =0.238, R= 0.873, p=0.001), but there was higher betadispersion in the Disturbance treatment for the Active communities (Comparing Total v. Active for Disturbance mesocosms, Kruskal Wallis p=0.029). This suggests that there was Active community variability masked by the contributions of dead and dormant taxa to the Total community.

**Figure 4.**
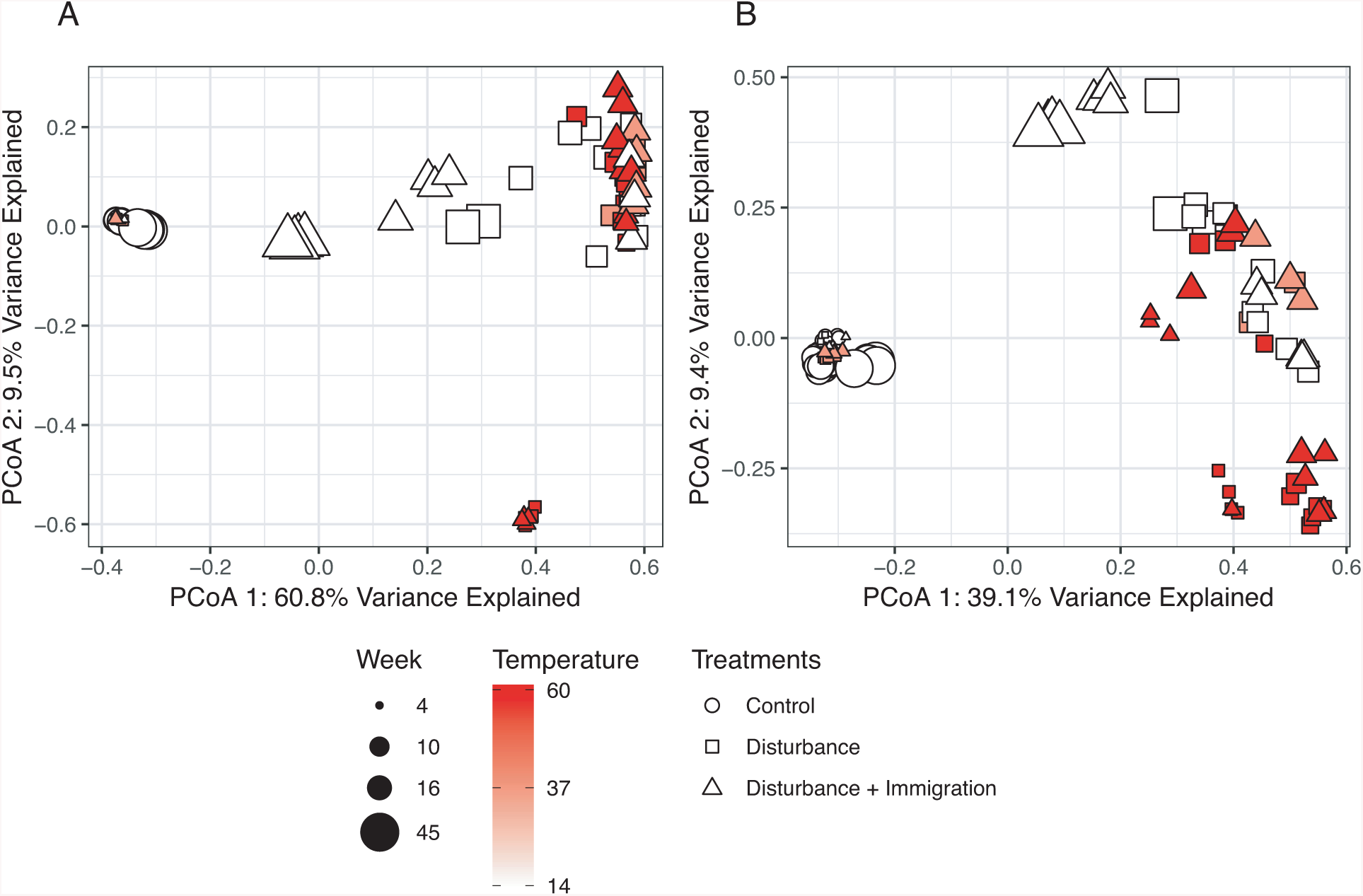
Changes in beta diversity over the disturbance experiment. Pairwise differences in community structure was quantified using pairwise Bray-Curtis dissimilarity and then ordinated using Principal Coordinates Analysis (PCoA). Time is shown by symbol size, and mesocosm temperature is indicated by heat colors, with the brightest red indicating the warmest time point. Control mesocosms are circles, Disturbance are squares, and Disturbance + Immigration are triangles. (A) PCoA of the Total community, assessed using sequencing of the 16S rRNA gene. (B) PCoA of the Active community, including only OTUs that had 16S rRNA:rRNA gene > 1.

**Figure 5.**
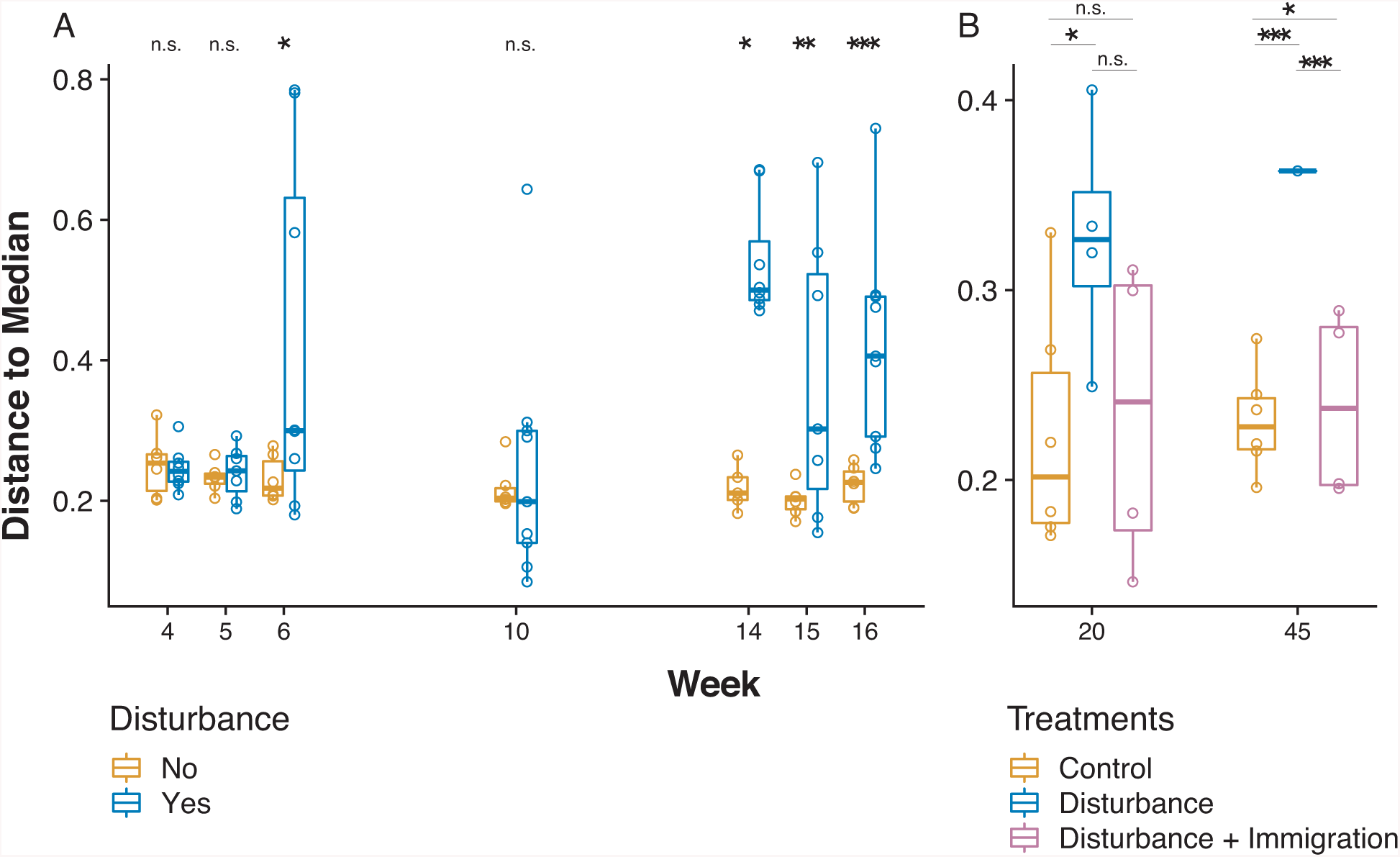
Changes in beta dispersion over the disturbance experiment. Beta dispersion, an indicator of variability in community structure, was quantified using the distance to the median in ordination space (Figure 4), which was constructed based on Bray-Curtis dissimilarity. (A) Changes in beta dispersion in Control (gold) and Disturbance (blue) mesocosms over the thermal press (weeks 4-16). (B) Changes in beta dispersion in Control, Disturbance, and Disturbance + Immigration (pink) mesocosms over the recovery period, weeks 20-45. The Disturbance + Immigration mesocosms received a dispersal event at week 17. Asterisks indicate significant differences with a Tukey’s Honestly Significant Difference post-hoc test (n.s. = not significant, * p<0.1, ** p<0.01, *** p<0.001). Note differences in y-axis ranges between A and B.

Replicate disturbed mesocosms (n=9, inclusive of Disturbance and Disturbance + Immigration) had highly reproducible responses to the press. They had high overlap in membership and overall synchronous trajectories, even after the immigration event at week 16 (33 of 36 PROTEST all R > 0.89 and false-discovery rate adjusted p-values < 0.05).

### Resistance and resilience

Using the Active community, we calculated resistance and resilience of the Disturbance treatment relative to the Control using community divergence from the first sampling time(Week4) as the reference (**Figure 6A**). Even in the Control communities, there was an initial drop in similarity between weeks 4 and 5, which we attribute to the bottle effect. However, after that the Control communities remain relatively stable with no additional divergence, while the Disturbance communities decrease to their maximum divergence at week 10 (60°C).

**Figure 6.**
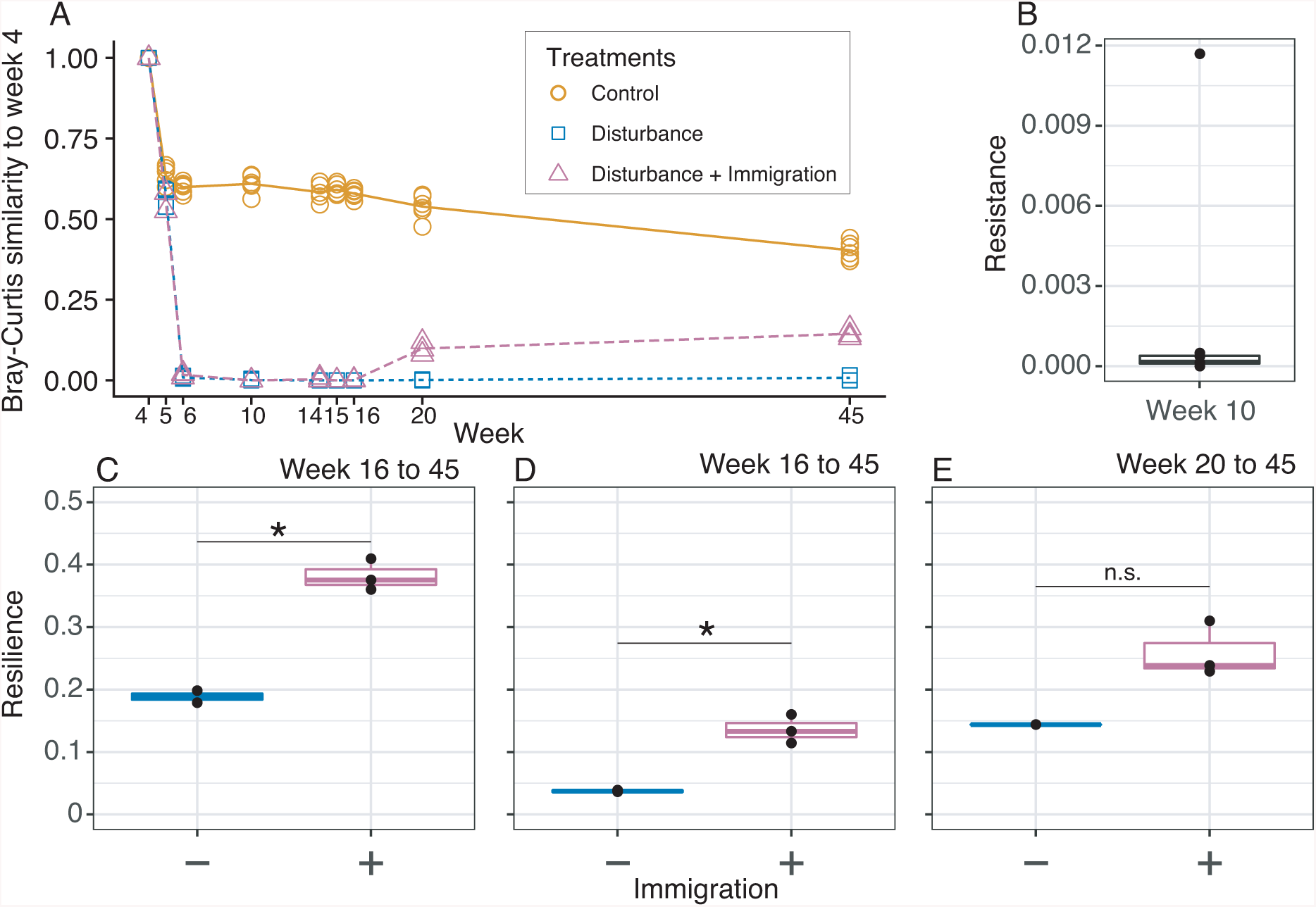
Resistance and resilience of soil mesocosm communities to a thermal press. (A) Temporal series of community divergence from pre-disturbance community (week 4) in Control (gold solid line), Disturbance (blue short dashed line), and Disturbance + Immigration (pink long dashed line) to calculate resistance and resilience. (B) Resistance of disturbed mesocosms at week 10, the time point of maximum community change after the thermal press begins. (C-E) Resilience of disturbed mesocosms without (-) and with (+) immigration, calculated after the thermal press is released (week 16) for the (C) full recovery to week 45, (D) initial recovery to week 20, and also for (E) long-term recovery from weeks 20 to 45. Asterisks indicate significant differences by a Kruskal Wallis test (n.s. = not significant, * p<0.1).

Disturbed communities with Immigration converge slightly after the dispersal event. Overall resistance was low (**Figure 6B**), and resilience reached its maximum, 0.41, in the immigration treatment between weeks 16 (the time point at which the thermal press was released) and the final week 45, but ranged from a minimum of 0.04 between week 16 and 20 in the Disturbance without immigration treatment (**Figure 6C-E**). Immigration enhanced resilience from week 16 to week 20 (Kruskal Wallis p value 0.034) and from week 16 to week 45 (Kruskal Wallis p value 0.083), but not from week 20 to 45, possibly because of insufficient power (Kruskal Wallis p value 0.180). There were only two Disturbance replicates (out of five) that met our rarefaction threshold for week 45.

For the recovery period (weeks 16-45), we wanted to assess the relative contributions of activity dynamics and immigration to the overall beta diversity in **Figure 2A**. We calculated the relative contribution of activity dynamics by identifying taxa that switched from an active or inactive state to the other during this recovery period. We found that these dynamically active taxa contributed 11.7% to 58.9% (median 28.9%) of the observed beta diversity, while immigrants contributed 8.1% to 27.3% (median 15.5%) of the observed beta diversity during the same time period.

### Activity dynamics of responsive taxa

To understand potential roles of dormancy initiation and resuscitation in driving community resistance and resilience, we wanted to distinguish taxa that changed in their activity or their detection over the course of the disturbance. Taxa that fell below detection (there was no rRNA gene detected in a particular sample) were distinguished from taxa that became inactive (rRNA:rRNA gene shifted from > 1 to < 1). For this analysis, we used the Active community but coded taxa that fell below detection as NAs to distinguish them from inactive taxa, which were coded as 0. Notably, taxa that fell below detection could have been either active, inactive, or locally extinct.

To conservatively attribute activity dynamics, we restricted this analysis only to the taxa that were among the 50 most abundant over the course of the experiment (**Figure 7A**). Within this set, we detected no purely resistant taxa that were consistently active throughout the experiment. This finding agrees with the analyses showing low resistance (**Figure 6B**) and substantial shifts in the Disturbance communities (**Figure 5**).

**Figure 7.**
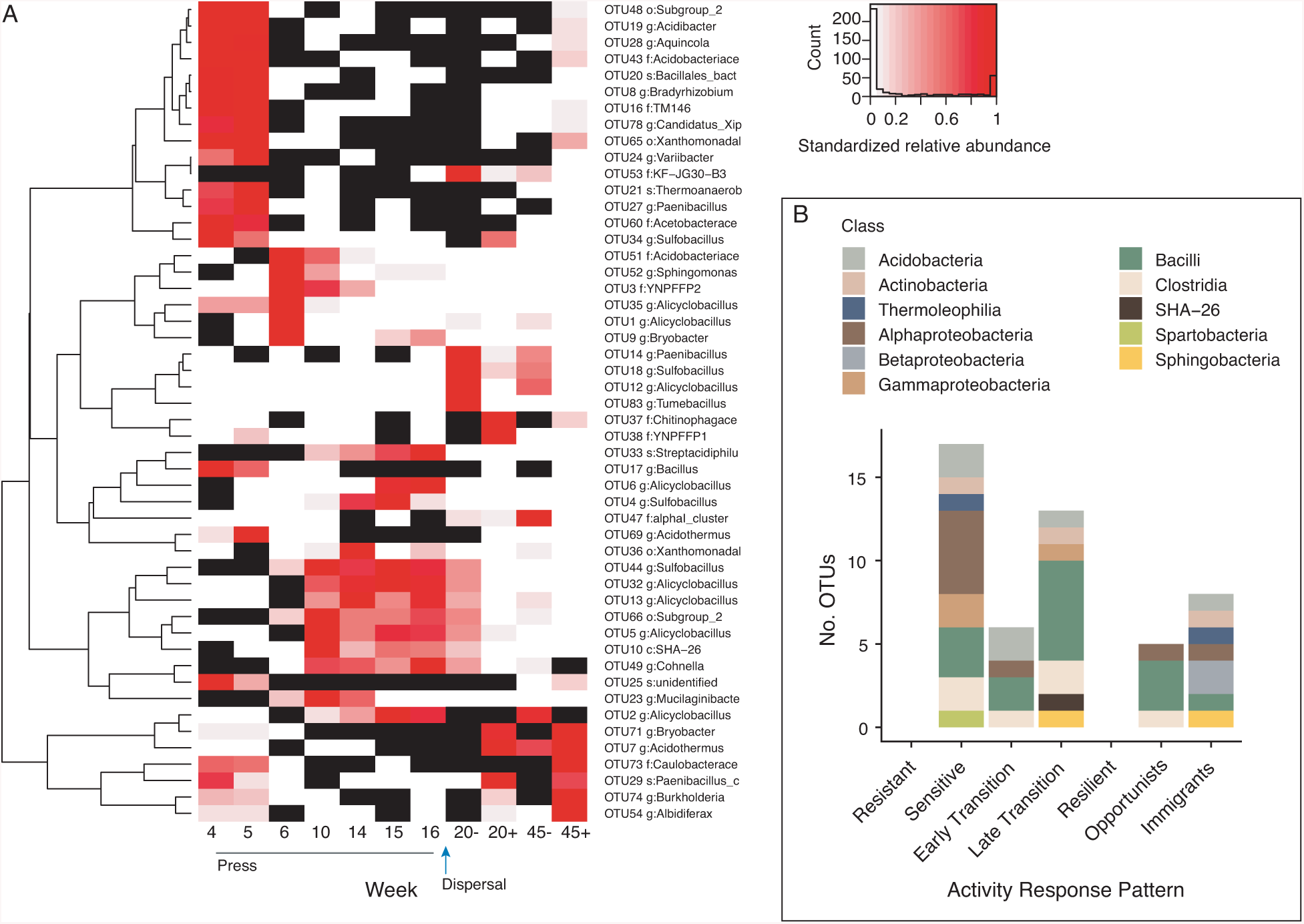
The activity dynamics of the 50 most abundant taxa in response to the press disturbance. (A) Heatmap and dendrogram of abundant taxa reveal common patterns of detection and activity. Black cells are taxa that were undetected (coded as NA) in the 16S rRNA gene (DNA) community, and white cells are taxa that were detected in the DNA but had 16S rRNA:rRNA gene < 1 (inactive, coded as 0). The heat gradient indicates each taxon’s abundance relative to its maximum observed in disturbance treated mesocosms during the experiment. Immigration is indicated for weeks 20 and 45 by minus (no) and plus (yes) signs. (B) Summary of activity response patterns to the disturbance of the top 50 taxa, including resistant, sensitive, early and late transition, resilient, opportunist, and immigrant taxa.

We detected 17 taxa that were sensitive to the disturbance (**Figure 7B**). Sensitive taxa were active prior to the warming but became inactive or dropped below detection during the warming, and then did not reactivate. We also detected 19 transition taxa that were inactive prior to the warming, active during the warming, and then became inactive after the stressor was released. Because there was no external dispersal into the system, these thermotolerant taxa were likely in the dormant pool of the soil. We could divide these responses generally into early and late transition taxa. There were 6 early transition taxa that became active during week 5 or 6 of the experiment, but then became inactive at weeks 10 and 14. There were also 13 late transition taxa that remained inactive during weeks 5 and 6 but became active during weeks 10 and 14.

Among the top 50 taxa, we did not detect purely resilient taxa that were active prior to the warming, became inactive during the warming, but then reactivated after the return to ambient temperature. This suggests that dormancy strategies responsive to warming were not a substantial contributor to member preservation, nor to eventual re-seeding. Instead, opportunists and immigrants facilitated resilience in the mesocosms. Five opportunists were inactive or below detection prior to and during the warming, but then activated after the temperature returned, likely due to resuscitation. Eight immigrants were generally active prior to the warming, dropped to below detection or became inactive during the warming, and then in the end, were active again only in the Disturbance + Immigration treatment (and not in the Disturbance mesocosms without immigration).

### Relationships between taxon activity and abundance

The conventional thought is that relative abundance is the outcome of growth and therefore an indicator of fitness, and so high relative abundance is indicative of recent or current activity in the environment. However, we detected a weak, but statistically supported, inverse (log10) relationship between OTU 16S rRNA:rRNA gene ratio and relative abundance for those taxa with an rRNA:rRNA gene ratio >1 (**Figure 8A**, Pearson’s R = -.14, p < 0.0001). This result is in agreement with other studies that have suggested that rare taxa may have high activity levels relative to their abundance in the community [42–46]. We present it here to be transparent that there are likely additional active but rare members that contribute to stability that have not been considered in our analyses.

**Figure 8.**
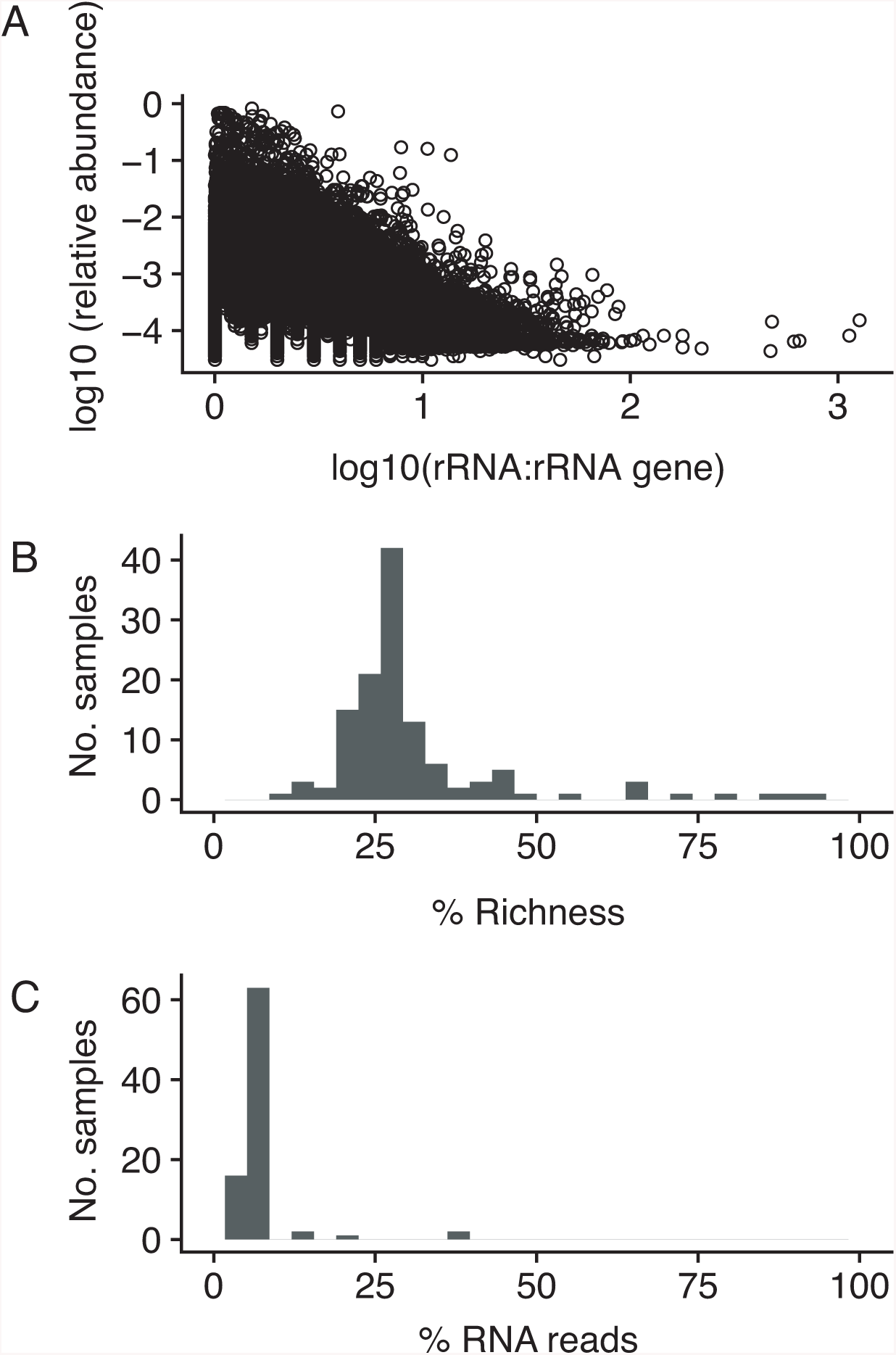
Taxon activity and abundance relationships. (A) Log10 relative abundance and log10 rRNA:rRNA gene ratio were inversely correlated. Each point is a different OTU detected in the dataset that had 16S rRNA:rRNA gene greater than or equal to 1. (B) Distribution of percent sample richness (No. OTUs detected, inclusive of DNA and RNA datasets) that were phantom taxa (16S rRNA detected but not 16S rRNA gene). (C) Distribution of percent RNA reads attributed to phantom taxa.

The inverse relationship between activity and abundance could not include taxa that had RNA but no DNA detected (aka “phantom taxa”, [44]) because they have an undefined 16S rRNA:rRNA gene ratio. We make clear that, to be conservative, phantom taxa (that have RNA but no DNA detected) were not included in the analyses, and that rare taxa that had high activity ratios were not included in the description of activity response patterns among the top 50 most abundant taxa. On balance, phantom taxa contributed proportionally few rRNA reads and few unique OTUs to the dataset (**Figure 8B and 8C**). However, there were a few exceptions, including five samples that had >10% rRNA reads and > 50% of richness attributed to phantom taxa. Four of these were from the Disturbance mesocosms at week 14 (peak-thermal press), and one sample was from week 16, at the end of the press. These samples also had relatively low richness and community size (**Figure 2** and **3**). We speculate that, by reducing community size and likely also total microbial biomass, the disturbance indirectly provoked relatively higher contributions by phantom taxa and conditionally rare taxa [47].

## Discussion

These results show that both dispersal and local dormancy dynamics, including activation and inactivation, can contribute to overarching patterns of community resilience. The dispersal event simulated in this experiment posed an optimistic scenario: well-mixed, control soils were mixed into disturbed soils to maximize the volume of the disturbed soil that came into contact with the inoculum. Regardless, by all metrics (beta diversity, alpha diversity, community size), immigration was very successful. Our data directly show that dispersal can augment resilience towards recovery. Given that the influences of dispersal on community assembly has been investigated previously (often indirectly for bacterial and archaeal microbiomes, as inferred from the contributions of stochastic or neutral processes e.g., [19,48–51]), this result is in agreement with the consensus of the literature that dispersal and dispersal limitation can matter for assembly [52–54].

A new result is that local resuscitation also contributes to microbiome community transitions during disturbance, and to resilience after the stress is released. Among the most abundant taxa, there were near equal numbers of taxa that contributed to resilience via opportunistic resuscitation and to resilience via immigration. Therefore, both mechanisms – local resuscitation and regional immigration – are important for microbiome stability. The microbial dormant pool is important for maintaining microbial diversity [43] and has evolutionary implications for traits that persist within inactive populations [55]. To make more explicit the role of dormancy dynamics for community disturbance responses (e.g., [56]), the phenomenon of the “storage effect” underpins modern coexistence theory [57] and refers to the ability of competing species to coexist when their growth and activities are separately partitioned over time, typically in dynamic environments [58]. Given the severity of the thermal stressor in Centralia and in this experiment, our results suggest that the soil microbial dormant pool is deep, in that it contains functionality for distinctive conditions, like thermal stress, that are not within the expected range of environmental variability.

Another goal of the experiment was to understand the reproducibility of member resuscitation given the press, and from the same soil. Because we observed high divergence in the hot field soil communities in Centralia that was not attributable to any measured soil environmental variable, including temperature, we hypothesized that stochastic resuscitation from the soil could initiate priority effects (e.g., [9]), leading to divergent hot communities. However, there was strong reproducibility among replicate disturbed mesocosms, suggesting that there were particular microorganisms that consistently responded to the thermal stress from the same soil. Therefore, we interpret that resuscitation in response to the thermal stress was largely deterministic, and that observed divergences among hot soil communities in the field may be instead attributed to either differences local edaphic factors that were unmeasured, or different underlying dormant pools, or stochasticity in regional dispersal. Thus, our hypothesis regarding priority effects by stochastic resuscitation was not supported.

Moving forward, there are several insights gleaned from this experiment. For soil, measuring dispersal in the field is difficult, given the various means by which microorganisms may arrive to a locality, including wind, ground water, and invertebrate vectors. However, there are several methods, each with their own caveats and biases [18,35,59,60], to measure activity and the pool of dormant organisms, and so this has become possible even with field soils. We recommend to collect member activity data for soils, to characterize the dormant pool, and to develop database infrastructure to support meta-analyses of these distinct activity-linked data resources. Also, microbiome stability is a dynamic process that involves both transition and resilience, and longitudinal series that are inclusive of the entire trajectory are informative. Characterizing the full disturbance trajectory will allow for quantification of the changing mechanisms that support stability, and will facilitate prediction of microbiome outcomes to new stressors. In our experiment, one week of stress was sufficient to observe community sensitivity (by week 5, the control and the disturbance treatments were statistically different), but 29 weeks after the stress was not sufficient to observe complete recovery, though it seems that recovery is possible given the trajectory toward the controls. For many soils, we expect that this time frame of response may be typical [61] and it can be used to inform future studies.

To conclude, this experiment shows both dispersal and dormancy dynamics can contribute to soil microbiome resilience in response to a press stress. Specifically, resuscitation of thermotolerant members contributed to transition during press, and then immigration provided a substantial boost to recovery beyond what was achieved with resuscitated opportunists. Because activity responses to the disturbance were consistent, these results suggest that predictive insights into microbiome resilience can be advanced more generally. We expect that accounting for mechanisms of local resuscitation and regional dispersal together will advance quantitative understanding of environmental microbiome stability.

## Acknowledgements

This work was supported by the National Science Foundation under Grant No DEB#1749544. This work was supported in part by Michigan State University through computational resources provided by the Institute for Cyber-Enabled Research. We thank Johnathon Higgins for technical assistance in the laboratory.

